# Machine Vision–Enabled Lateral Flow Immunoassay Using Functionalized Gold Nanoparticles for Point-of-Care Cardiac Biomarker Detection

**DOI:** 10.1101/2025.10.22.683875

**Authors:** Yuhan Lu, Xiaoyun Peng, Zhao Yin, Xiaoqin Fan, Jingchun Fan, Yali Mi, Guangchao Li

## Abstract

Acute myocardial infarction (AMI) remains one of the most prevalent and fatal cardiovascular disease. Given the critical diagnostic significance of cardiac troponin I (cTnI) and myoglobin (Myo) in AMI, there is an urgent clinical demand for rapid and accurate detection methods to improve patient outcomes and reduce mortality. To meet this need, we developed a sensing platform that integrates functionalized gold nanoparticle-based lateral flow immunoassay (AuNPs-LFIA) with a machine vision model, enabling rapid quantitative detection of cTnI and Myo. By covalently conjugating antibodies to gold nanoparticles (AuNPs@Ab), we achieved greater probe specificity and stability. The integration of machine vision algorithms allowed quantitative readouts within 8 minutes, a 46.7% improvement of the detection time compared to conventional methods (15 minutes). The platform achieved limits of detection of 0.224 ng/mL for Myo and 0.071 ng/mL for cTnI, with excellent correlation to commercial kits (R^2^ > 0.99). Overall, these results demonstrate that machine vision-enhanced AuNPs-LFIA offers an efficient, sensitive and reliable strategy for point-of-care testing (POCT) in cardiovascular diagnostics, particularly in resource-limited settings.

## 1. Introduction

Achieving high sensitivity and rapid diagnosis remains a major challenge in point-of-care testing (POCT). Existing POCT approaches primarily fall into two categories: immunoassay-based and molecular detection-based methods. Although molecular diagnostics offer high accuracy, they often involve extensive sample preparation and prolonged assay times of 3-4 hours^[1, 2]^. Immunoassays. Likewise, immunoassays such as enzyme-linked immunosorbent assay (ELISA) require 3-5 hours to complete^[3-5]^, making both approaches impractical for urgent, near-patient application. In contrast, paper-based lateral flow immunoassays (LFIA) deliver qualitative results within approximately 15 minutes, offering a substantial time advantage^[6, 7]^. However, in emergency medicine, particularly for acute myocardial infarction (AMI)-a prevalent and highly fatal cardiovascular disease-clinicians demand both rapid and quantitatively reliable readouts^[8, 9]^. Early AMI diagnosis further benefits from the simultaneous detection of multiple cardiac biomarkers, with myoglobin (Myo) and cardiac troponin I (cTnI) recognized as critical indicators^[10-14]^. Their complementary sensitivity and specificity provide a stronger basis for clinical decision-making.

Commonly employed LFIA probes include fluorescent microspheres, quantum dots, and colloidal gold (AuNPs). While fluorescent probes can enhance sensitivity, they are prone to photobleaching and require specialized detection equipment, increasing cost and compromising portability-limitations that hinder their use in resource-limited or decentralized POCT settings^[15-18]^. By contrast, AuNPs are widely used due to their intuitive colorimetric readout and operational simplicity. Yet, conventional AuNP-based probes face three primary limitations: (i) antibody (Ab) is typically adsorbed via noncovalent interactions, reducing probe stability and specificity; (ii) results rely on visual interpretation, introducing subjectivity; and (iii) quantitative accuracy remains limited^[19-21]^. These shortcomings highlight the need for functionalized probes with enhanced stability and specificity, coupled with rapid and reliable quantitative readout strategies.

Recent advances in machine vision offer a promising solution for rapid, quantitative LFIA^[22-27]^. Unlike traditional hardware-based optimization, machine vision leverages image recognition and pattern learning to accelerate detection, improve accuracy, and enhance robustness^[28-30]^. Motivated by this potential, we developed a functionalized gold nanoparticle-based lateral flow immunoassay (AuNPs-LFIA) integrated with a deep learning-based machine vision framework for the rapid and quantitative detection of Myo and cTnI. By covalently conjugating antibodies to synthesized functional AuNPs, probe stability and specificity were significantly improved. The integration of machine vision reduced assay time from 15 minutes in conventional LFIA to 8 minutes while enabling automated quantitative readout, minimizing subjective errors and improving accuracy and reproducibility (Figure 1). Collectively, this work establishes a rapid and practical strategy for the early detection of myocardial injury and demonstrates the broader potential of combining nanotechnology with deep learning for point-of-care quantitative diagnostics.

**Figure 1.**
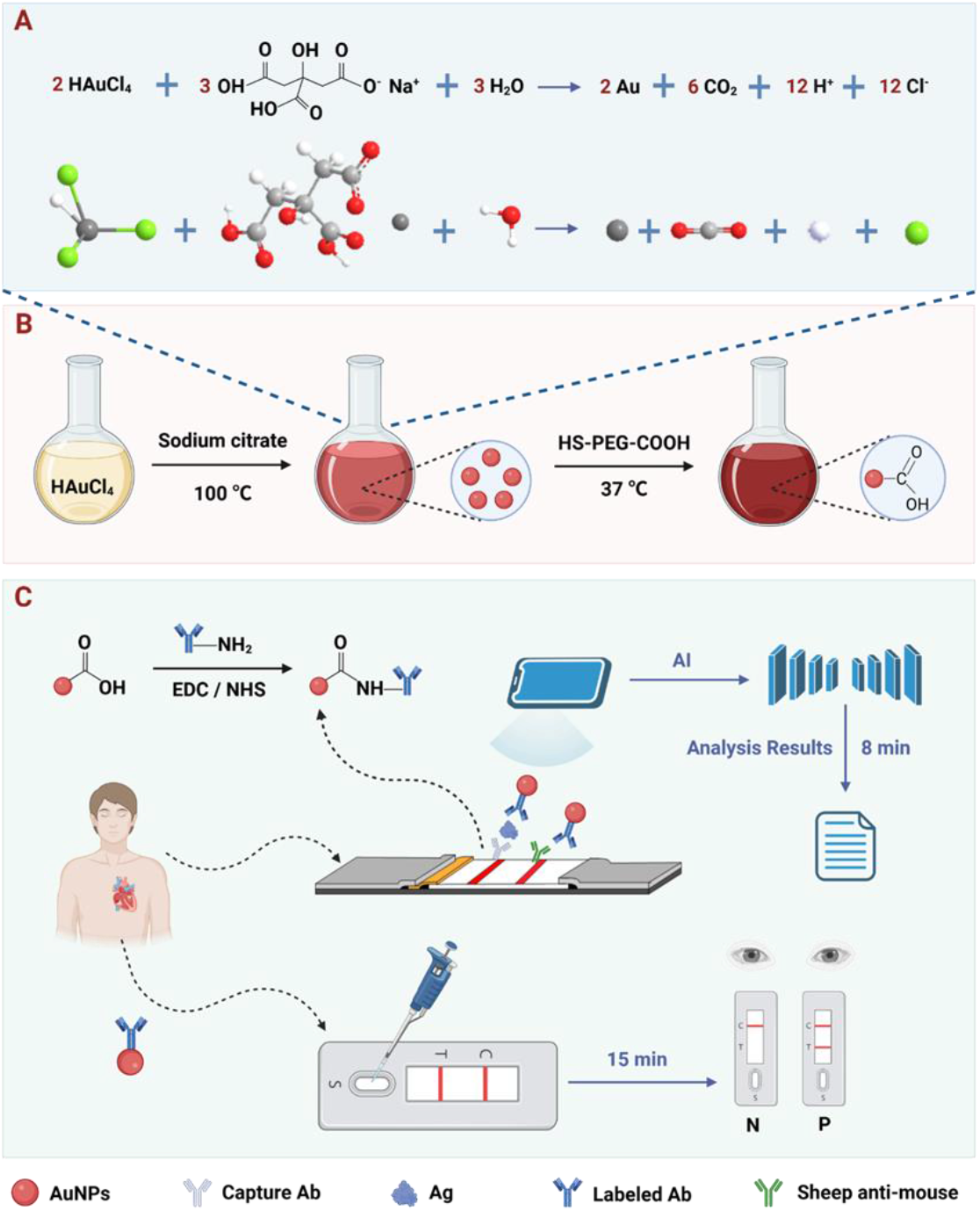
Schematic illustration of the workflow for AMI biomarker detection. (A) Principle of AuNP synthesis using chloroauric acid and sodium citrate. (B) Schematic representation of AuNP synthesis. (C) Comparison of the AuNP-supported LFA with AI-assisted analysis and commercial LFAs for **AMI** biomarker detection.

## 2. Results and Discussion

### 2.1. Characterization of AuNPs and AuNPs@Ab

To prepare functionalized gold nanoparticles (AuNPs) covalently conjugated with antibodies (AuNPs@Ab) for colorimetric detection, AuNPs were synthesized via the classical sodium citrate reduction method. In this approach, chloroauric acid (HAuCl_4_) served as the gold precursor, and sodium citrate acted as both reducing and stabilizing agent. Under heating conditions, Au^3+^ ions were reduced to elemental gold, yielding monodisperse AuNPs with uniform size (**Figure 1A**). To introduce carboxyl functional groups on the AuNP surface for subsequent covalent Ab conjugation, the AuNPs were modified with thiol-terminated polyethylene glycol with a carboxyl group (HS-PEG-COOH). This modification relies on the strong covalent Au-S bond between the thiol group and the gold surface, ensuring stable PEG attachment while presenting terminal carboxyl groups as active sites for Ab conjugation (Figure 1B).

Following surface modification, the carboxyl groups were activated with 1-ethyl-3-(3-dimethylaminopropyl) carbodiimide (EDC) and N-hydroxysuccinimide (NHS), facilitating amide bond formation between the activated carboxyl groups and primary amines of lysine residues in the target antibody. This reaction resulted in the covalent immobilization of antibodies on the AuNP surface (**Figure 2A**). To characterize the morphology, size, and surface properties of the synthesized AuNPs and AuNPs@Ab probes, multiple analytical techniques were employed, including transmission electron microscopy (TEM), dynamic light scattering (DLS), zeta potential measurements, and UV-vis spectroscopy. TEM imaging revealed that the synthesized AuNPs were highly dispersed spherical particles with an average diameter of approximately 20 nm (Figure 2B), indicating uniform size distribution and favorable dispersibility, advantageous for stable migration through LFIA strips. DLS analysis showed a hydrodynamic diameter of 31.25 ± 0.46 nm with a narrow size distribution, confirming monodispersity in aqueous solution (Figure 2C). Upon Ab conjugation, the zeta potential shifted significantly from −24.0 ± 0.53 mV to +16.6 ± 0.55 mV (Figure 2D), indicating a substantial alteration in surface charge and successful covalent attachment of antibodies to the AuNP surface. Moreover, the UV-vis absorption peak of AuNPs exhibited a slight red shift from 524 nm to 527 nm after Ab conjugation, further corroborating effective antibody immobilization (Figure 2E).

**Figure 2.**
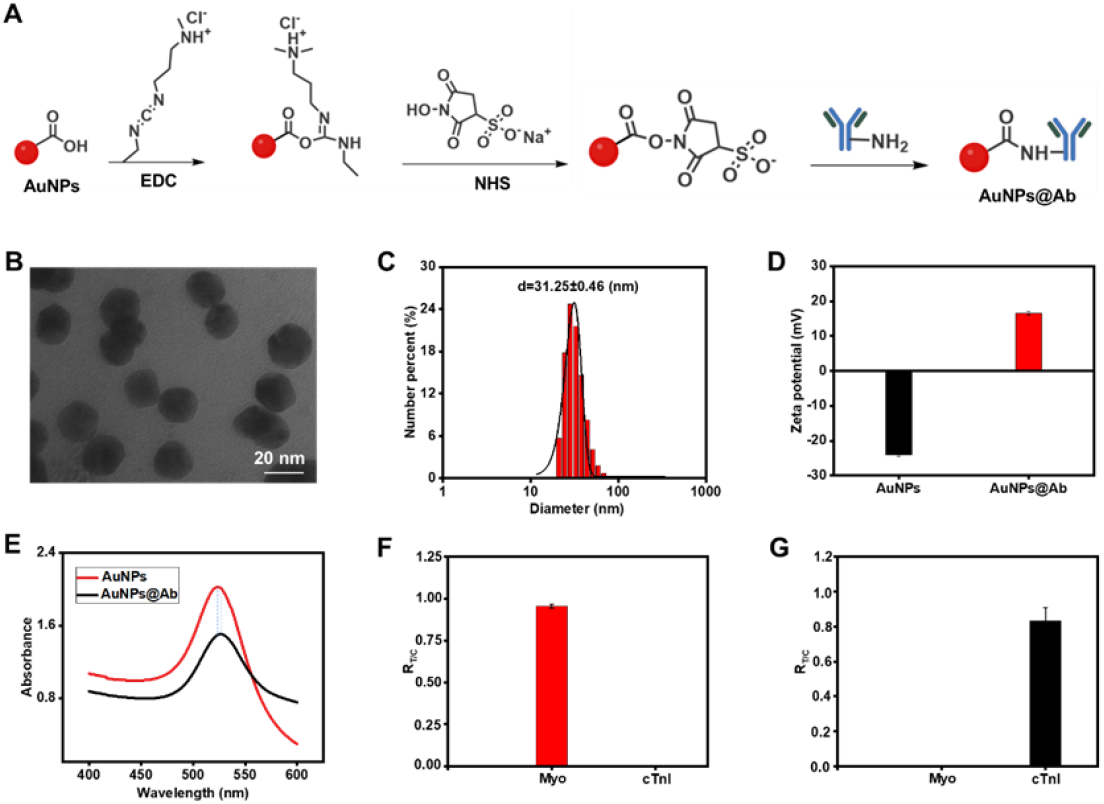
Characterization of AuNPs and AuNPs@Ab. (A) Schematic diagram of the process for the covalent coupling Ab of AuNPs. (B) TEM characterization of AuNPs. (C) Dynamic light scattering characterization of AuNPs. (D) Zeta sites characterize AuNPs and AuNPs@Ab. (E) Ultraviolet absorption spectra of AuNPs and AuNPs@Ab. (F) Specificity analysis of Myo. (G) Specificity analysis of cTnI.

To further evaluate the specificity, cross-reactivity assays were performed using AuNPs functionalized with monoclonal antibodies against either Myo or cTnI. As shown in Figures 2F and 2G, no detectable colorimetric signal was observed at the test line (T line) when Myo antibody-conjugated AuNPs were challenged with cTnI antigen, or when cTnI antibody-conjugated AuNPs were exposed to Myo antigen. These findings demonstrate the absence of cross-reactivity between the two probes, confirming that the selected antibodies possess high specificity and selectivity for their respective target antigens, thereby minimizing the likelihood of false-positive results and nonspecific interactions.

### 2.2. Performance analysis based on gold nanoparticles

LFIA is a portable diagnostic platform that enables rapid target detection based on the principle of capillary action. It is mainly composed of five key components: a sample pad, a conjugate pad, a nitrocellulose (NC) membrane, an absorbent pad, and a poly(vinyl chloride) (PVC) backing card. The sample pad serves to receive the test sample, while the conjugate pad is preloaded with AuNPs@Ab probes. The NC membrane is marked with T line and C line for target identification and assay validation. The absorbent pad maintains the capillary flow of the liquid (**Figure 3A**). To achieve optimal detection sensitivity and ensure consistent performance in practical applications, we first systematically optimized the detection conditions. Variables included the volume of AuNPs@Ab probes, the applied sample amount, and the concentration of capture antibodies on the NC membrane. Under a fixed antigen concentration, the detection performance was evaluated by comparing the signal intensity ratio of the T and C lines (T/C). Taking Myo as a model analyte, the optimal conditions were determined to be 1 μL AuNPs@Ab, 60 μL sample volume, and a capture antibody concentration of 1 mg/mL, whereas for cTnI, the optimal conditions were 1 μL AuNPs@Ab, 50 μL sample volume, and 0.8 mg/mL capture antibody (Figure S1-S2).

**Figure 3.**
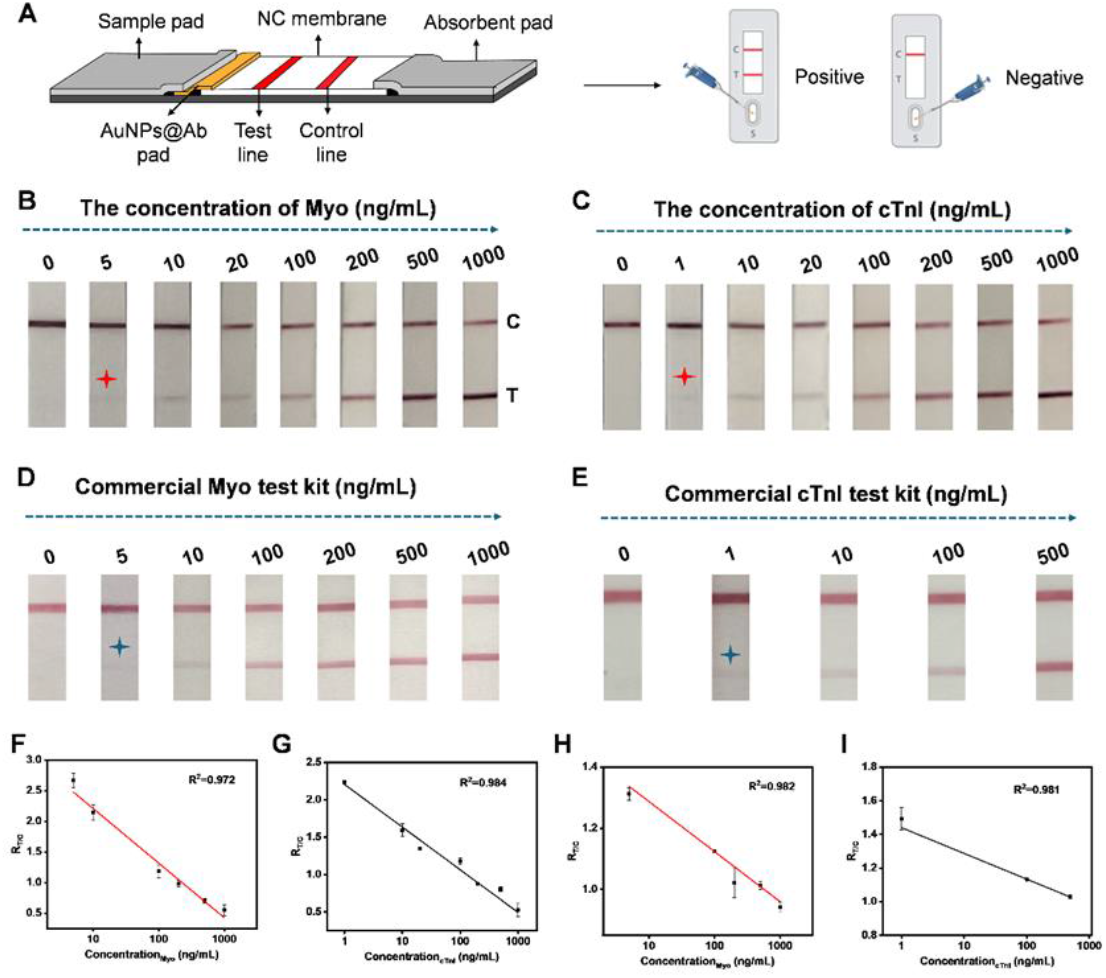
Detection of Myo and cTnI using AuNP-based LFA. (A) Schematic illustration of the LFA configuration and representative detection outcomes. (B) Readout results of AuNP-based LFA for varying concentrations of Myo. (C) Readout results of AuNP-based LFA for varying concentrations of cTnI. (D) Readout results of a commercial LFA for different concentrations of Myo. (E) Readout results of a commercial LFA for different concentrations of cTnI. (F) Calibration curve showing the linear relationship between Myo concentration and corresponding signal intensity.(G) Calibration curve showing the linear relationship between cTnI concentration and corresponding signal intensity. (H) Calibration curve of Myo obtained from the commercial LFA. (I) Calibration curve of cTnI obtained from the commercial LFA. Data are presented as mean ± SD (n = 3).

Under optimized detection conditions, we further evaluated the performance of the platform for both biomarkers. For Myo, a concentration gradient ranging from 5 to 1000 ng/mL was tested, and the colorimetric response at the T line was monitored. As shown in Figure 3B, the T line colorimetric intensity gradually increased with rising Myo concentrations, demonstrating a clear concentration-dependent response. Quantitative analysis was performed using ImageJ to extract the R-channel value of both the T and C lines, followed by calculation of the T/C ratio (R_T/C_) and regression fitting against concentration. The calibration plot revealed a strong linear relationship within the tested range (Figure S3A and Figure 3F), described by the regression equation y = -0.892 Lg(x) + 3.10 (R^2^ = 0.972). The visual limit of detection was determined to be 5 ng/mL. Similarly, for cTnI, a concentration range of 1-1000 ng/mL was investigated. As shown in Figure 3C, the R-channel value at the T line decreased with increasing cTnI concentrations. Quantitative analysis was performed using ImageJ to extract the R-channel value of both the T and C lines, followed by calculation of the T/C ratio (R_T/C_) and curve fitting against concentration. The calibration plot revealed a strong linear correlation between signal intensity and analyte concentration (Figure S3B and Figure 3G), described by the regression equation y 127 = -0.570 Lg(x) + 2.21 (R^2^ = 0.984). The visual limit of detection was determined to be 1 ng/mL.

To further compare the performance of functionalized AuNPs-LFIA with that of commercial AuNPs-LFIA, we tested Myo and cTnI using commercially available LFA strips. In the commercial LFAs, the detection signals (R-channel) of Myo and cTnI decreased with increasing concentrations (Figure 3D-E). The linear regression equations of y = -0.164 Lg(x) + 1.45 (R2 = 0.982) and y = -0.153 Lg(x) + 1.44 (R^2^ = 0.981), respectively, indicating a good linear correlation between signal intensity and analyte concentration (Figure S4, Figure 3H-I). As shown by comparing Figure 3B with Figure 3D, the variation in R_T_ values observed in functionalized AuNPs-LFIA was markedly greater than that of the commercial counterparts. These findings indicate that antibody-functionalized AuNPs substantially enhance the specific binding efficiency to the target, thereby improving colorimetric signal sensitivity and resolution.

### 2.3. Machine vision-based analysis of LFIA

To establish a high-precision and rapid-response LFIA detection system, we developed and deployed a deep learning-based machine vision model for real-time quantitative analysis of cardiac biomarkers cTnI and Myo. As illustrated in **Figure 4A**, the workflow was implemented using the YOLOv11 object detection framework. The training dataset was derived from video frames captured during experimental assays. After YOLOv11 training, the optimized model was obtained and capable of simultaneously detecting the T and C lines on multiple LFIA and reliably extracting their RGB channel values. In this study, the red (R) channel was selected as the key detection metric to enhance color contrast and quantitative sensitivity. The trained model was integrated into an embedded lower-level system, where a CCD industrial camera periodically captured strip images, which were then processed by the model in real time to automatically generate T/C ratios for user interpretation.

**Figure 4.**
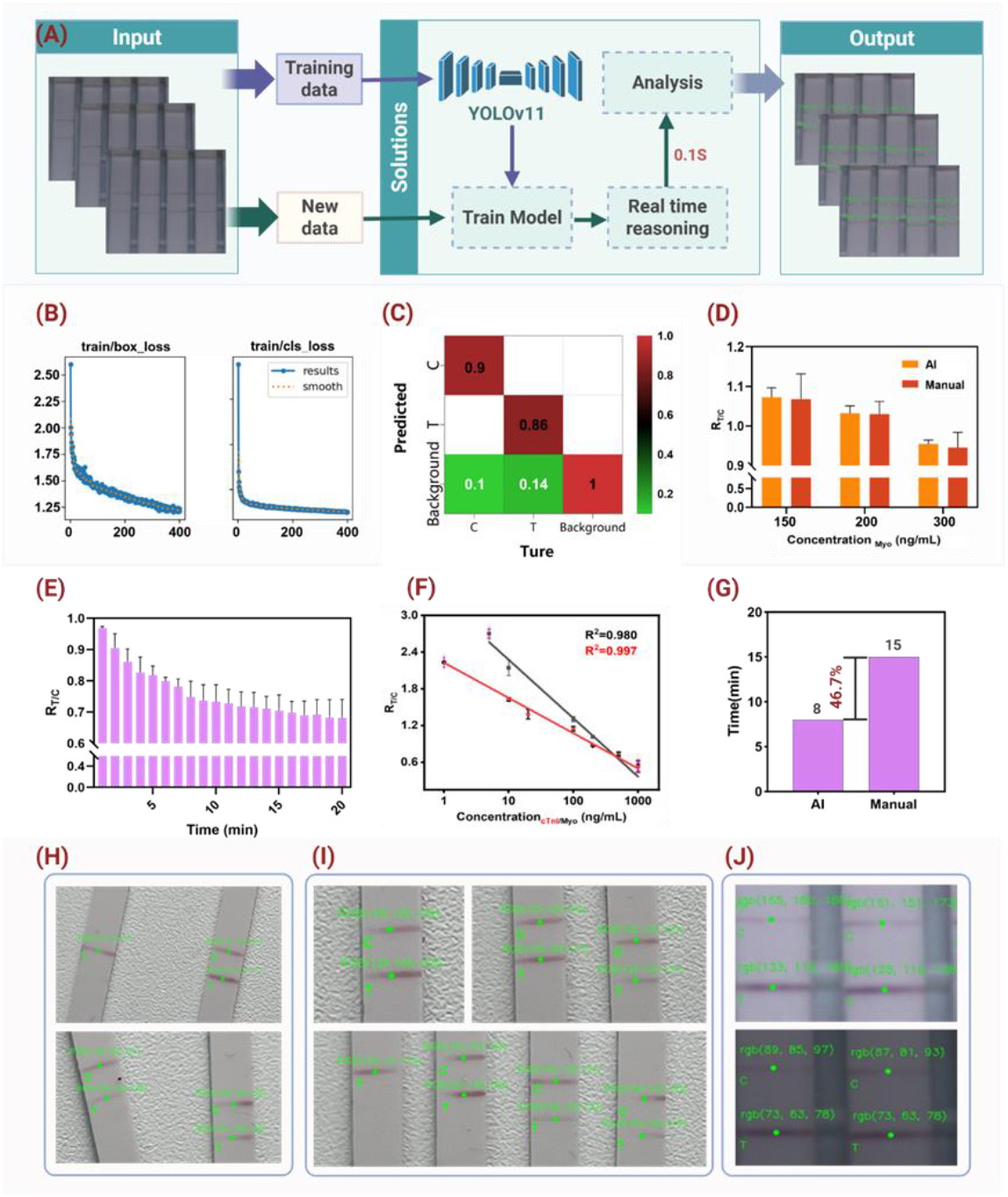
Machine vision-based analysis of LFIA. (A) Schematic workflow of machine vision-based detection of the T and C lines on LFIA. (B) Loss curves during the training of the machine vision model, including bounding box localization loss and classification loss. (C) Normalized confusion matrix of the model’s classification results, showing the accuracy of identifying different regions. (D) Statistical comparison of T/C ratios in the R channel obtained by machine vision versus manual analysis at Myo concentrations of 150, 200, and 300 ng/mL. (E) Dynamic change of the R_T/C_ value over time at a Myo concentration of 300 ng/mL. (F) Standard curves for cTnI and Myo generated based on machine vision detection results. (G) Comparison of analysis time between machine vision and manual detection, with the former saving approximately 7 min per test. (H-J) Model generalizability tests under varying positions, angles, quantities, and ambient lighting conditions.

During training, both the bounding box loss and classification loss steadily decreased and converged around the 400th epoch, indicating robust model performance (Figure 4B, Figure S5). Figure 4C presents the model’s classification performance for C- and T-line recognition (Figure S6-S7). The normalized confusion matrix shows an accuracy of 90% for C-line detection and 86% for T-line detection, with minimal misclassification of background regions. Most errors mainly occurred at the early color-development stage of T and C lines, when the strip coloration was not fully developed.

To assess detection accuracy, Myo concentrations of 150, 200, and 300 ng/mL were measured in triplicate by both machine vision detection and manual colorimetric measurement. Results from the two approaches were highly consistent (Figure 4D), with an average agreement of 99.16%. The maximum error was 1.38% for machine vision and 4.63% for manual detection, indicating that machine vision reduced errors by approximately 3.35-fold and substantially improved the quantitative analysis reliability.

Furthermore, to assess dynamic detection performance, the R-channel T/C ratio was monitored over time at a Myo concentration of 300 ng/mL (Figure 4E). The T/C ratio gradually decreased with reaction time, reaching a plateau at ∼8 minutes and stabilizing by 15 minutes. Thus, the 8-minute time point was selected as the final detection node to balance sensitivity and efficiency.

Figure 4F shows the standard calibration curves for cTnI and Myo obtained via machine vision. The cTnI linear equation was y = -0.955Lg(x) + 3.23 with R^2^ = 0.997, y = -0.577Lg(x) + 2.23 with R^2^ = 0.980, demonstrating strong linear responses and validating the method’s applicability across multiple targets. The limits of detection (LOD) for Myo and cTnI were calculated using the formula LOD = 3δ/S, where δ represents the standard deviation of the low-concentration group and S denotes the slope of the calibration curve, yielding values of 0.224 ng/mL and 0.071 ng/mL, respectively. A comparison of detection time between machine vision and manual measurement (Figure 4G) showed that automated detection reduced the average assay time by 7 minutes, improving efficiency by ∼46.7%, making it highly suitable for point-of-care diagnostics and large-scale screening.

Finally, to evaluate model generalizability, tests were conducted under varying strip positions, orientations, quantities, and ambient lighting conditions. As shown in Figures 4H-J, the system maintained robust performance under these complex conditions, underscoring its strong adaptability and broad applicability of the proposed machine vision model.

### 2.4. Validation of the Reliability of the AuNPs-LFIA Platform

To further evaluate the reliability and practical applicability of the AuNPs-LFIA platform integrated with the YOLOv11 deep learning algorithm for clinical sample analysis, a series of spiked experiments were designed to simulate real clinical conditions. Specifically, Myo and cTnI, two key biomarkers of myocardial injury, were spiked into FBS to generate a set of test samples. These samples were then analyzed using both the developed AuNPs-LFIA platform and commercial assay kits, and the detection performance and consistency of the two methods were compared across different concentration levels. As shown in **Figures 5A** and 5C, the AuNPs-LFIA platform exhibited a clear concentration-dependent response for both Myo and cTnI, with R-channel signal intensity decreasing as analyte concentration increased. Notably, when combined with the YOLOv11 model, the LFA platform was capable of automatically identifying the T and C lines and accurately calculating the T/C ratio, significantly enhancing image recognition accuracy and data processing efficiency. Even at low analyte concentrations, the platform maintained high sensitivity and discriminative capability, a critical feature for the early detection of myocardial injury. To further assess the concordance between the LFA platform and commercialized LFIA, regression analysis was performed on the results obtained by the two methods, and correlation curves were plotted (Figures 5B and 5D). The Myo detection data showed an excellent fit with a correlation coefficient (R^2^) of 0.986, indicating high agreement between the methods; similarly, the R^2^ for cTnI was 0.996, confirming the platform’s strong consistency and broad applicability across different myocardial injury biomarkers. Beyond analytical performance, the AuNPs-LFIA combined with YOLOv11 offered significant advantages in workflow simplicity, response speed, and operational convenience. The lateral flow design enabled rapid antigen-antibody interaction and signal generation with only a single droplet of sample applied to the sample pad, completing the entire assay within 8 min. Simultaneously, the YOLOv11 algorithm provided real-time image recognition and quantitative analysis of the T line, eliminating subjective errors associated with manual interpretation and markedly improving result objectivity and reproducibility.

**Figure 5.**
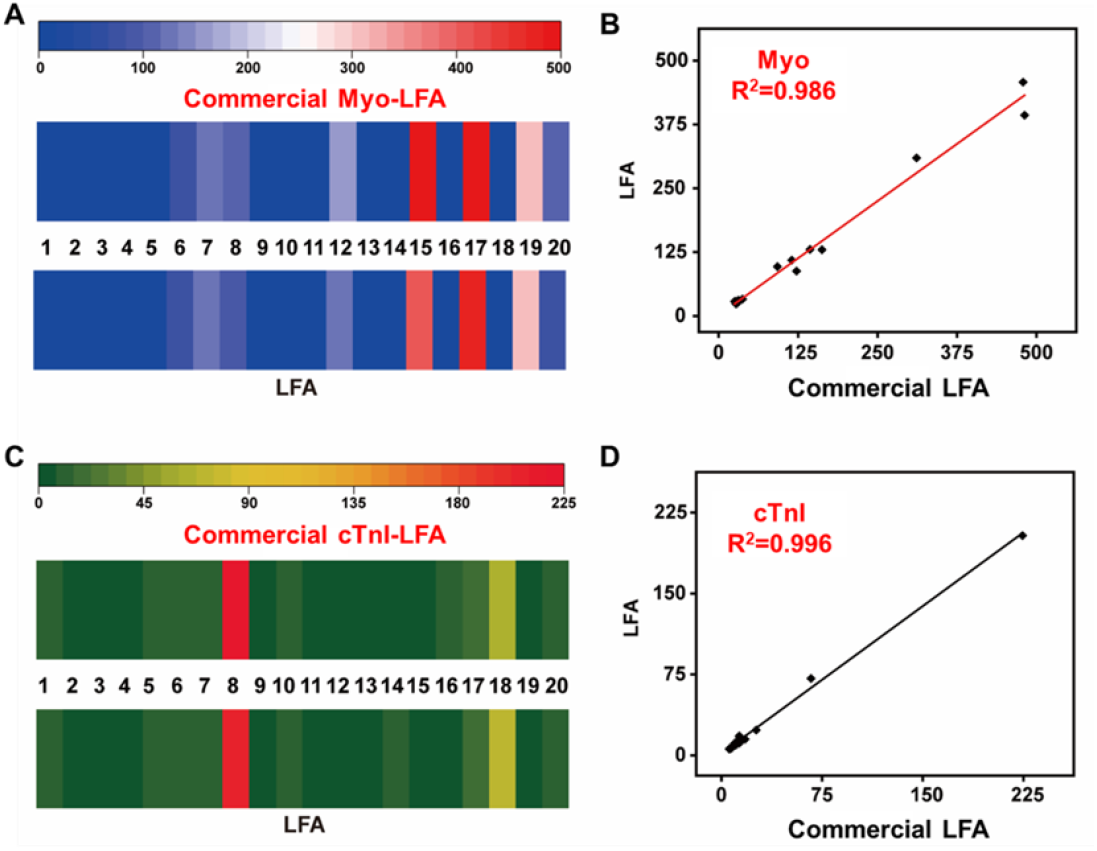
Comparative analysis of spiked Myo and cTnI samples using the AuNPs-LFIA platform and commercial ELISA kits. (A) Comparison of Myo detection results at different spiked concentrations, showing good agreement between the AuNPs-LFIA platform and commercial LFIA. (B) Correlation analysis between Myo concentrations measured by the AuNPs-LFIA platform and commercial LFIA, with a correlation coefficient of R^2^ = 0.986. (C) Comparison of cTnI detection results in spiked samples between the AuNPs-LFIA platform and commercial LFA. (D) Correlation analysis of cTnI detection results between the AuNPs-LFIA platform and commercial LFA, demonstrating a high degree of consistency.

## 3. Materials and Methods

### 3.1. Materials and reagents

Chloroauric acid hydrated (HAuCl_4_) was from Beijing Xinsheng Baitai Technology Co., LTD (Shanghai, China). Sodium citrate, bovine serum albumin (BSA), fetal bovine serum (FBS), sucrose and sodium chloride were from Shanghai Maclean Biochemical Technology Co., LTD (Maclin, Shanghai). Thiol-polyethylene glycol-carboxylic acid was from Guangzhou Jiexin Biotechnology Co., LTD. Nitrocellulose membranes, glass fiber and absorbent pad were from Shanghai Jiening Biotechnology Co., LTD (Shanghai, China). Myo 212 (capture Ab), Myo 213 (labeled Ab), Myo antigen, cTnI monoclonal antibody (capture Ab), cTnI monoclonal antibody (labeled Ab), cTnI antigen and goat anti-mouse IgG (H+L) were from Qianyuan Shunze Technology Co., LTD (Qingdao, China). The kits of Myo and cTnI were from Taobao.

### 3.2. Synthesis of gold nanoparticles

Gold nanoparticles (AuNPs) with an average diameter of approximately 20 nm were synthesized using the classical citrate reduction method^[31]^. Briefly, 13 μL of 1 mM chloroauric acid (HAuCl_4_) solution was added to 50 mL of ultrapure water in a 100 mL round-bottom flask and brought to a boil under magnetic stirring. In parallel, a 1% (w/v) sodium citrate solution was prepared by dissolving 0.03 g of sodium citrate in 3 mL of ultrapure water. Once the HAuCl_4_ solution reached boiling, 2 mL of the sodium citrate solution was rapidly added. During the reaction, the color of the solution changed sequentially from pale yellow to gray, purple, and finally to wine red, indicating the successful formation of AuNPs. The mixture was maintained at boiling for an additional 15 minutes and then allowed to cool naturally to room temperature. The resulting AuNPs were stored at 4 °C for further use. To improve the colloidal stability and introduce functional groups, the surface of the AuNPs was modified with thiol-terminated polyethylene glycol bearing a carboxyl group (HS-PEG-COOH). A 5 mg/mL HS-PEG-COOH aqueous solution was prepared, and 200 μL of this solution was added to 25 mL of the synthesized AuNPs. The mixture was incubated on a shaker (320 rpm) at room temperature for 2 h. Subsequently, the solution was left undisturbed at 4 °C overnight to promote the formation of stable Au-S bonds between the thiol groups and the AuNP surfaces. Unbound HS-PEG-COOH molecules were removed by repeated washing with ultrapure water, and the carboxyl-functionalized AuNPs were finally resuspended in 25 mL of ultrapure water and stored at 4 °C for future use.

### 3.3. Covalent conjugation of antibodies to gold nanoparticles

Carboxyl-functionalized AuNPs were conjugated with antibodies via a carbodiimide-mediated amide coupling strategy. Briefly, 1 mL of carboxylated AuNP solution was incubated with 10 μL of 0.4 mM EDC (1-ethyl-3-(3-dimethylaminopropyl) carbodiimide hydrochloride) and 0.1 mM NHS (N-hydroxysuccinimide) in MES buffer (pH 6.0) at room temperature with gentle shaking for 15 min to activate the surface carboxyl groups into reactive NHS ester intermediates. Following activation, the mixture was centrifuged at 10000 rpm for 6 min to remove excess EDC and NHS, and the pellet was resuspended in ultrapure water. Next, 15 μg of antibody was added to the activated AuNPs and incubated at room temperature for 1 h to facilitate covalent coupling between the NHS esters and the primary amine groups on the antibody via stable amide bond formation. Afterward, 100 μL of 10% (w/v) bovine serum albumin (BSA) was added and incubated for an additional 30 min to block non-specific binding sites. Finally, the conjugates were centrifuged at 12000 rpm for 6 min, and the supernatant was discarded. The resulting AuNPs@Ab conjugates were resuspended in 50 μL of running buffer for storage and future use.

### 3.4. Preparation of LFA test strips

The lateral flow immunoassay (LFA) strip was composed of five components: a sample pad, a nitrocellulose (NC) membrane, an absorbent pad, a conjugate pad (glass fiber), and a polyvinyl chloride (PVC) backing card. Glass fiber was selected as the sample pad material and pretreated by immersing it in a borate buffer solution (pH 7.5) containing 5% bovine serum albumin (BSA), 2% sucrose, and 0.5% Tween-20, followed by air drying at room temperature. Subsequently, the sample pad, conjugate pad, NC membrane, and absorbent pad were sequentially assembled onto the PVC backing, with each pad overlapping the adjacent one by 3 mm. The capture antibodies for myoglobin (Myo) and cardiac troponin I (cTnI), as well as the goat anti-mouse IgG antibody (control line), were diluted in phosphate-buffered saline (PBS) to final concentrations of 0.8 mg/mL, 1 mg/mL, and 0.5 mg/mL, respectively. Using a dispenser, the antibodies were dispensed onto the NC membrane at a rate of 0.5 μL/cm to form the test lines (T-lines) and control lines (C-lines). After dispensing, the NC membrane was dried at 37 °C for 2 h. Finally, the assembled sheet was cut into 3 mm-wide individual strips using a cutter. The test strips were sealed in foil pouches containing desiccant and stored at room temperature until further use.

### 3.5. Optimization of immunoassay parameters for enhanced sensitivity

To achieve optimal detection sensitivity and ensure the reliability of the platform for practical applications, several critical parameters in the lateral flow immunoassay (LFIA) were systematically optimized. The parameters optimized included the volume of AuNPs@Ab conjugates, the sample volume, and the concentration of capture antibody at the test line. A one-variable-at-a-time strategy was applied, with the T/C intensity ratio at a fixed antigen concentration serving as the evaluation criterion for the comparative metric. Using myoglobin (Myo) as a model analyte, the results demonstrated that optimal performance was achieved with 1 μL of AuNPs@Ab, a 60 μL sample volume, and a capture antibody concentration of 1 mg/mL. For cardiac troponin I (cTnI), the optimal condition was obtained using 1 μL of AuNPs@Ab, a 50 μL sample volume, and a capture antibody concentration of 0.8 mg/mL.

### 3.6. AI Dataset Description

The dataset consists of 750 images of strips, exhibiting variations in strip position, number, orientation, and illumination, as well as diverse background conditions. Each image contains 1 to 4 strips, yielding a total of 4,500 annotated samples. The dataset was randomly divided into a training set (600 images), a validation set (100 images), and a test set (50 images). The training set was used for model optimization and weight learning, the validation set for hyperparameter tuning and model selection, and the test set for evaluating the model’s generalization performance on unseen data.

### 3.7. Data processing

Accuracy: The proportion of correctly predicted instances.

Formula: Accuracy=(TP+TN)/(FP+FN+TP+TN)

Recall (Sensitivity or True Positive Rate): The proportion of actual positives that are correctly identified.

Formula: Recall=TP/(FN+TP)

Precision (Positive Predictive Value): The proportion of predicted positives that are truly positive.

Formula: Precision=TP/(FP+TP)

### 3.8. Spiked Sample Analysis

To evaluate the quantitative performance of the constructed AuNPs-LFIA platform in practical samples, fetal bovine serum (FBS) was used as the matrix, and different concentrations of myoglobin (Myo) and cardiac troponin I (cTnI) were spiked to simulate clinical samples. Each spiked sample was tested in triplicate. For LFA detection, an image recognition approach incorporating the YOLO object detection algorithm was employed to capture and analyze the test strips. Images of the strips were acquired using a CCD, and the trained YOLO model automatically identified the test line (T line) and control line (C line), extracting their R-channel values to calculate the T/C ratio, which was then used for the quantitative determination of Myo or cTnI concentrations. All test strips were imaged under the same platform to ensure data consistency. For validation, commercial Myo and cTnI kits were used to quantify the concentrations of the respective biomarkers in the spiked FBS samples according the manufacturer’s protocols strictly.

## 4. Conclusions

In this study, we developed an AI-enhanced AuNPs-LFIA platform for rapid, sensitive and quantitative detection of cardiac injury biomarkers Myo and cTnI. AuNPs were functionalized with HS-PEG-COOH, introducing surface carboxyl groups that covalently bind to antibody amino groups via amide bonds, thereby endowing the probes with excellent specificity and stability. The performance of the AI-enhanced AuNPs-LFIA was evaluated against commercial kits using spiked serum samples, showing a high correlation between the two methods (R^2^ > 0.95) and confirming the analytical reliability of our platform. By training detection indices using machine vision, the quantitative analysis time was reduced from 15 min to 8 min while effectively eliminating the subjectivity associated with manual interpretation. This advancement significantly enhances the efficiency and accuracy of POCT, providing reliable clinical decision support for both healthcare professionals and non-specialist operators. Looking forward, this strategy has broad applicability and could be extended to the detection of other disease-related biomarkers, such as those associated with infectious diseases such as influenza, thereby facilitating rapid on-site diagnostics. As such, the AI-enhanced AuNPs-LFIA platform represents a practical and versatile platform for on-site diagnostics in diverse clinical and resource-limited settings.

## Supplementary Materials

The following supporting information can be downloaded at: https://www.mdpi.com/article/doi/s1, Figure S1: Condition optimization of Myo. Figure S2: Condition optimization of cTnI. Figure S3: Curve equation. Figure S4: The curve equation of commercial kits. Figure S5: Loss function. Figure S6: The relationhip between Recall and confidence level. Figure S7: The relationship between precision and recall.

## Author Contributions

The manuscript was written through contributions of all authors. **Yuhan Lu**: Writing – review & editing, Writing – original draft, Methodology, Investigation, Formal analysis. **Xiaoyun Peng**: Methodology. **Zhao Yin**: Methodology. **Xiaoqin Fan**: Methodology. **Jingchun Fan**: Data curation. **Yali Mi**: Data curation. **Guangchao Li**: Writing – review & editing, Supervision, Resources, Funding acquisition, Conceptualization. All authors have given approval to the final version of the manuscript.

## Funding

This work was supported by the **Young Scientists Fund of the National Natural Science Foundation of China** (Grant No. 82202034) and the **Xinjiang National Science Foundation Science and Technology Aid Project** (Grant No. 2022E02128). The authors gratefully acknowledge the financial support.

## Acknowledgments

We thank the Analysis and Testing Center of Jinan University for DLS, ζ-potential, TEM, and UV–Vis analyses, and Kexue Zhinnan for support with nanoparticle characterization.

## Data Availability Statement

The data presented in this study are available on request from the corresponding author.

## Conflicts of Interest

The authors declare no conflicts of interest.

## Notes

### Competing Interest Statement

The authors have declared no competing interest.

