## Supplemental Data 1 for "Machine Vision–Enabled Lateral Flow Immunoassay Using Functionalized Gold Nanoparticles for Point-of-Care Cardiac Biomarker Detection"

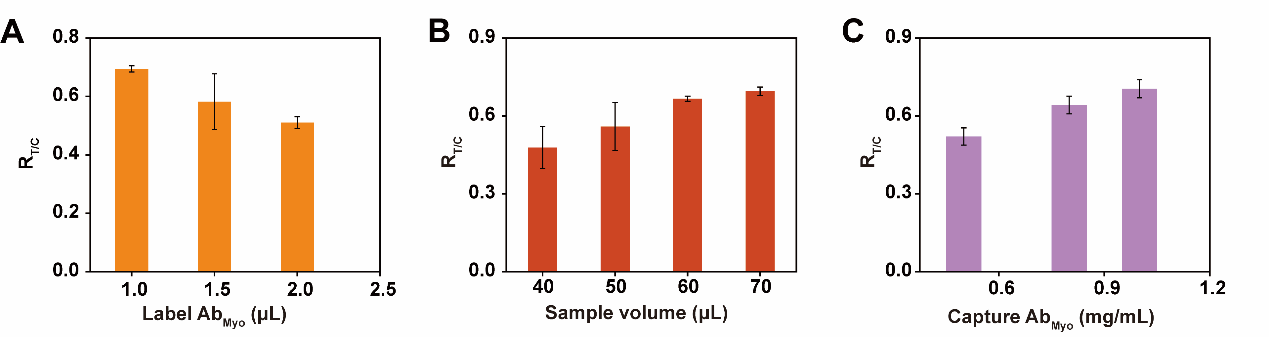

**Figure S1** Condition optimization of Myo. (A) Optimization of conditions for adding the volume of labeled Ab. (B) Optimization of conditions for adding sample volume. (C) Optimization of conditions for capturing antibody concentration.

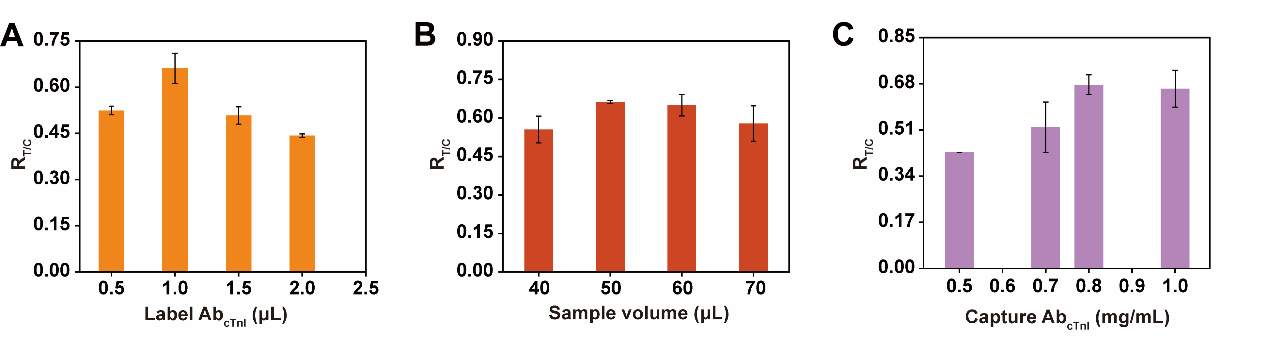

**Figure S2** Condition optimization of cTnI. (A) Optimization of conditions for adding the volume of labeled Ab. (B) Optimization of conditions for adding sample volume. (C) Optimization of conditions for capturing antibody concentration.

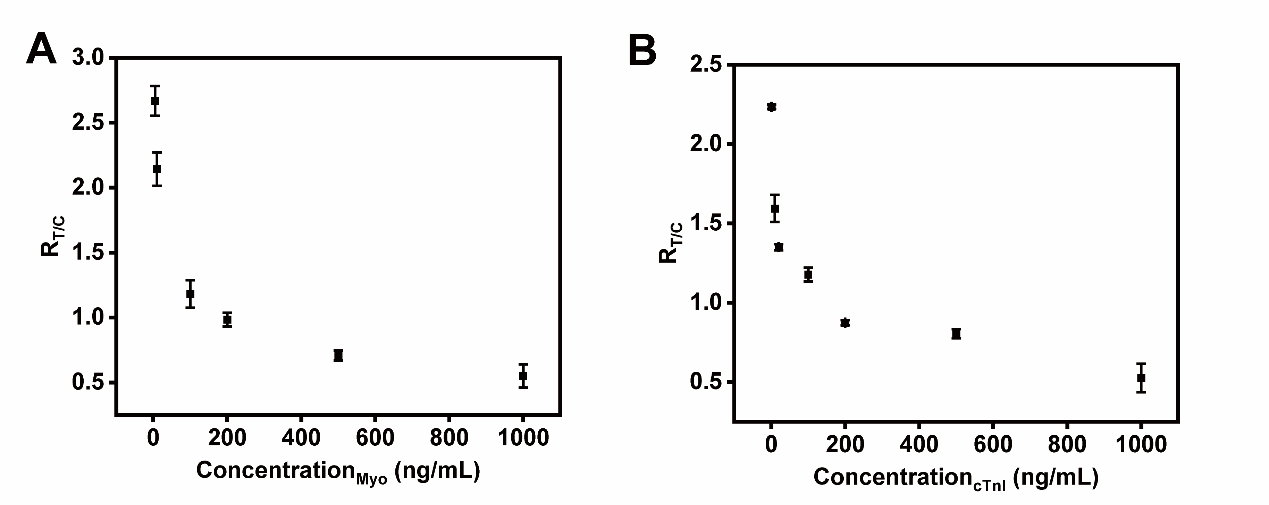

**Figure S3** Curve equation. (A) The curve equation of Myo. (B) The curve equation of cTnI.

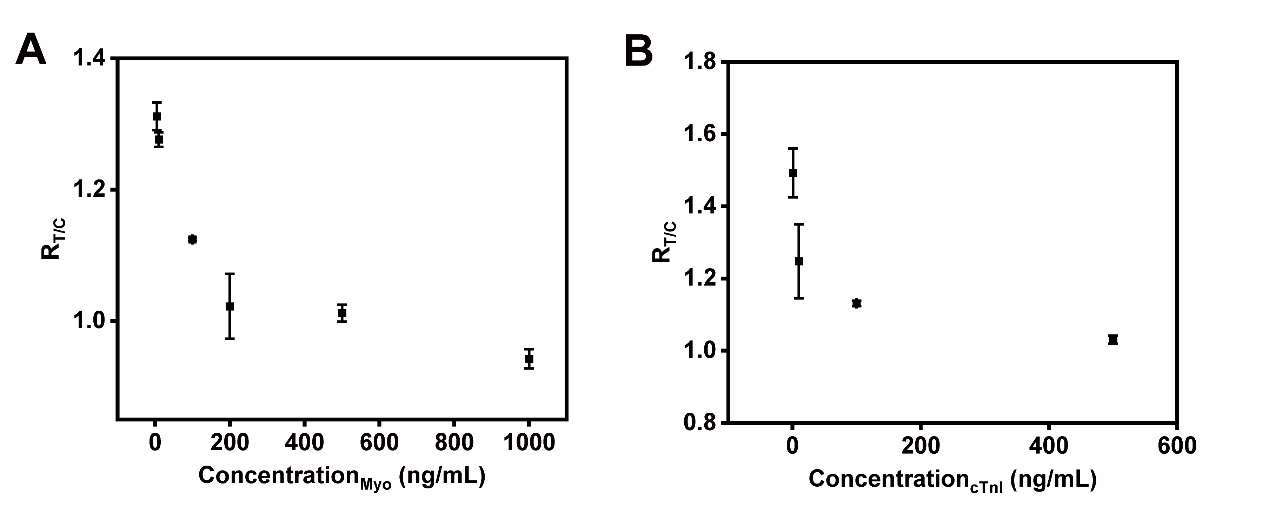

**Figure S4** The curve equation of commercial kits. (A) The curve equation of Myo. (B) The curve equation of cTnI.

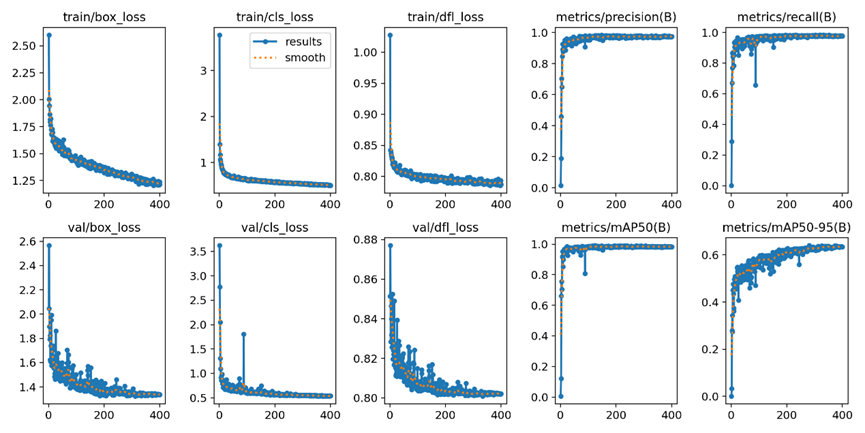

**Figure S5** Loss function. The loss function is used to measure the degree to which the predicted values of a model differ from the true values, and to a large extent, it determines the performance of the model.

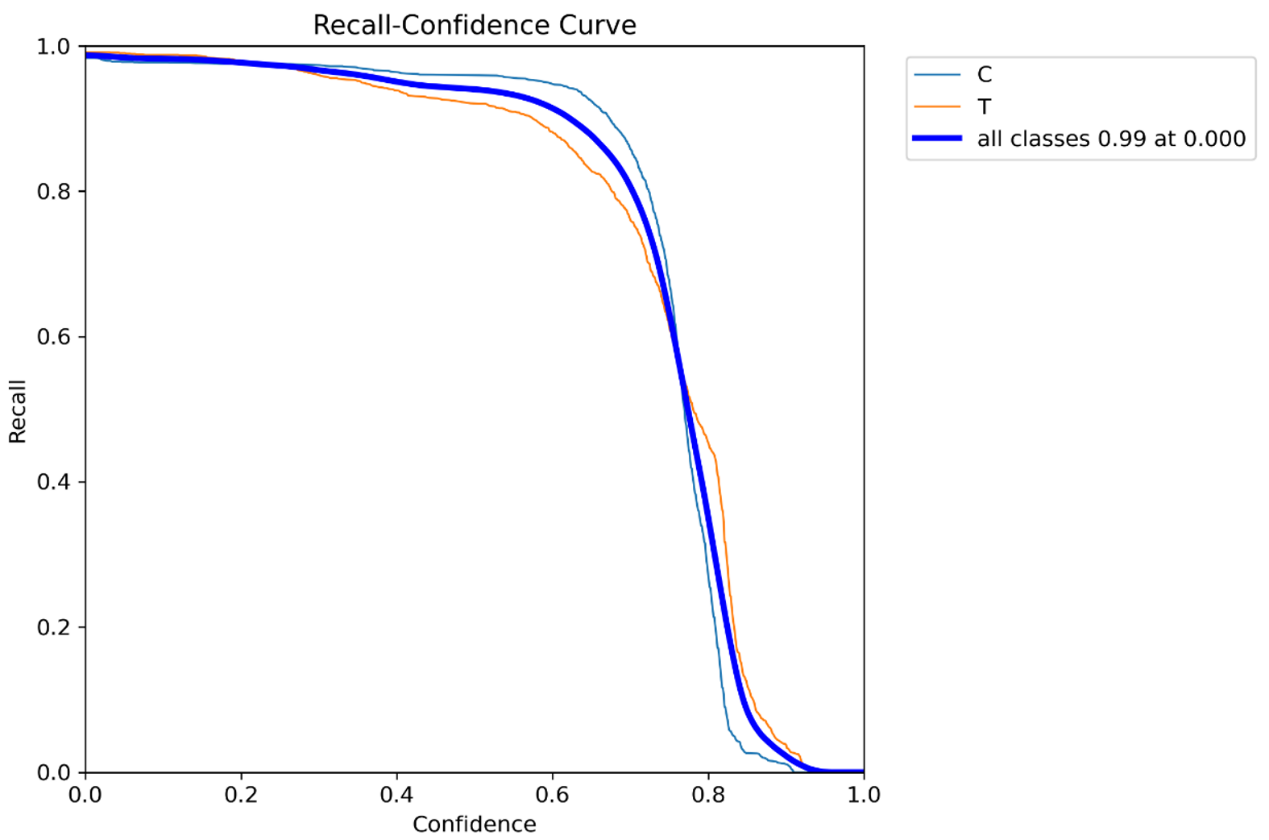

**Figure S6** The relationship between Recall and confidence level. When the confidence level is low, the detection is comprehensive but there are many misjudgments. When the confidence level is high, there are fewer misjudgments but more missed detections.

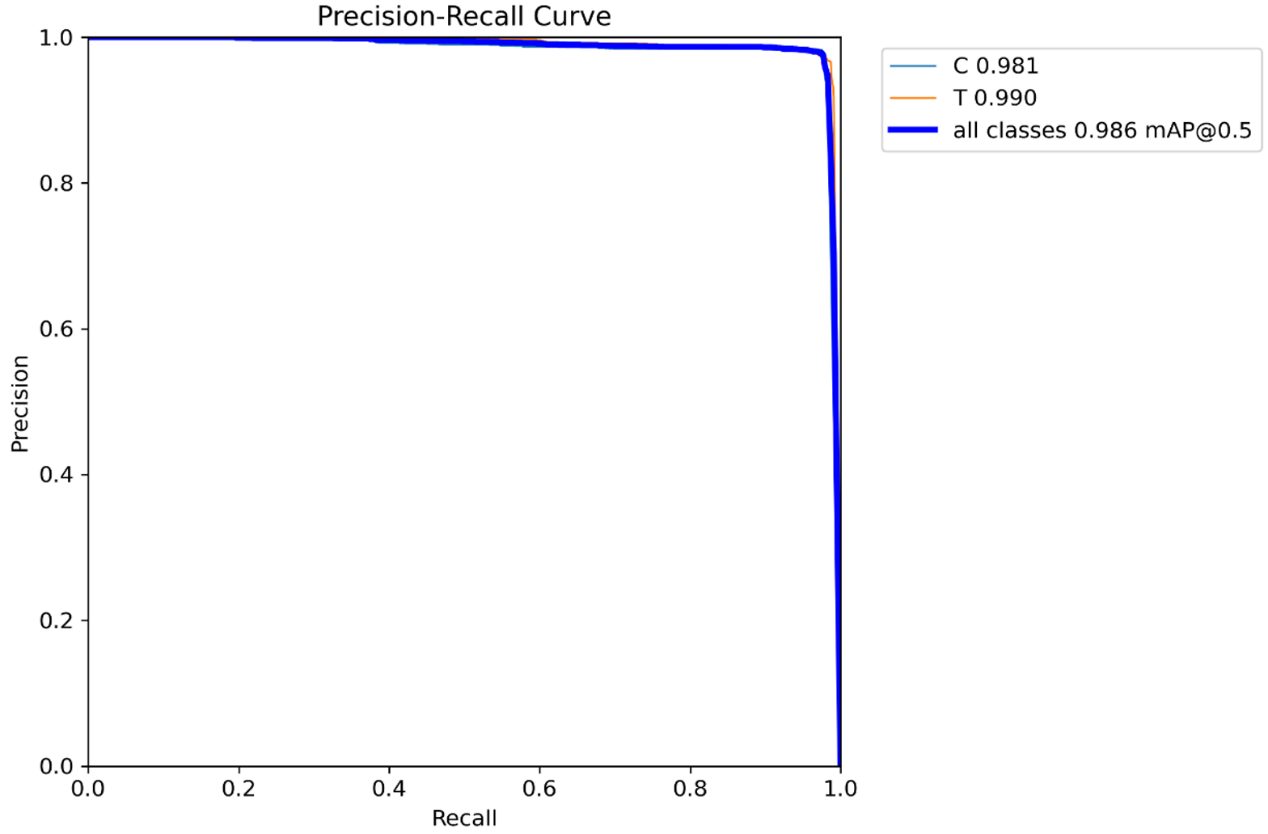

**Figure S7** The relationship between precision and recall. The larger the mAP (Mean Average Precision), the better the model performance. The area under the PR curve is AP, and the average value of AP for all categories is mAP.
